# Resistance-Breaking Root-Knot Nematodes from California Processing Tomato Fields Show Genomic Diversity and Variable Reproduction on Rotation Crops

**DOI:** 10.64898/2026.11.18.767369

**Authors:** Alison C. Blundell, Romnick Latina, Dadong Dai, Patricia Lazicki, Brenna Aegerter, Valerie M. Williamson, Shahid Siddique

## Abstract

Root-knot nematodes (RKNs) are a major threat for tomato growers worldwide. Management relies heavily on the plant resistance gene *Mi-1*, which has been bred into most processing tomato cultivars. However, repeated use of *Mi-1* has led to the emergence of resistance-breaking isolates throughout the world. Despite the growing impact of resistance-breaking RKNs, their frequency, distribution, and species identity across California remain poorly understood. In this study, we isolated RKN isolates from processing tomato fields at 14 sites across California, identified them to species, and evaluated their reproduction on current rotation crops to determine whether crop rotation could suppress these populations. All 14 sites yielded resistance-breaking populations, and most isolates were *M. incognita*. A SNP-based principal component analysis revealed substantial genetic diversity among several isolates, suggesting that resistance-breaking emerged independently in multiple fields rather than all spreading from a single population. In greenhouse assays, alfalfa displayed the lowest reproduction of both wild-type and resistance-breaking *M. incognita*, whereas reproduction on other rotation crops varied by isolate. Together, these results show that rotation decisions should be guided by local RKN population identity and host compatibility rather than generalized strategies across fields.

## Introduction

California has one of the most productive agriculture sectors in the United States, producing more than 400 commodities and supplying over one-third of the nation’s vegetables and more than three quarters of its fruit and nuts (California Department of Food and Agriculture, 2024). This productivity is supported by the state’s Mediterranean climate, which allows long growing seasons and cultivation of a diverse range of crops (Kauffman, 2003). Among the state’s top ten commodities, processing tomatoes rank eighth by value, generating roughly $1.5 billion in annual revenue (California Department of Food and Agriculture, 2023; USDA NASS, 2025).

Although California’s Mediterranean supports high crop productivity, it also favors many plant pathogens, including plant-parasitic nematodes. Among these, root-knot nematodes (RKNs; *Meloidogyne* spp.) are the most damaging, infecting over 3,000 plant species worldwide (Jones et al., 2013) and causing an estimated $8–10 billion in annual losses (Faske et al., 2026). The most damaging RKN species belong to the *Meloidogyne incognita* group (MIG), which includes *Meloidogyne incognita, M. javanica*, and *M. arenaria*, all of which occur in California. These nematodes penetrate roots and induce the formation of specialized feeding sites known as giant cells. They subsequently become immobile and depend on these cells for nutrition throughout the remainder of their lifecycle (Lin and Siddique, 2024). MIG RKNs reproduce asexually by mitotic parthenogenesis, with each female producing clonal progeny within a single egg mass containing 200 - 300 eggs (Triantaphyllou, 1985; Trudgill & Blok, 2001; Jones et al., 2013; Khan et al., 2023). In California’s processing tomatoes industry, RKNs account for an estimated $100 million in damage, although this figure likely underestimates their true economic impact, particularly given the limited management options available to growers (Koenning et al., 1999).

To combat MIG RKNs, the resistance gene *Mi-1.2* (hereafter referred to as *Mi-1*) has been incorporated into approximately 90% of commercial processing tomato cultivars (Smith, 1944; Gilbert and McGuire, 1956; Roberts, 1992; Williamson, 1998). In plants carrying *Mi-1*, nematodes can penetrate the roots but fail to establish feeding sites, ultimately leading to their death (Bleve-Zacheo et al., 1982; Paulson and Webster, 1972). However, studies have shown that *Mi-1* can lose efficacy when soil temperatures exceed 28^°^C. In addition, repeated cultivation of *Mi-1*-carrying cultivars, has imposed strong selection pressure on nematode populations (Djian-Caporalino et al., 2011; Dropkin, 1969; Abdul-Baki et al., 1996; Williamson, 1998; Verdejo-Lucas et al., 2013; de Carvalho, et al., 2015).

Concern over resistance-breaking RKNs has grown since their presence was first confirmed in the 1980s (Roberts et al., 1986). In addition to causing the direct yield losses, these nematodes create wounds that serve as entry sites for secondary pathogens such as *Fusarium noneumartii* (Fusarium stem rot and decline)*, Fusarium oxysporum (Fusarium wilt)*, *Sclerotium rolfsii* (southern blight), and *Pyrenochaeta lycopersici* (corky root) (Aegerter, 2025; Srinivas et al., 2019; Aycock, 1966; Blancard, 2012). Although root-knot nematodes require a living host to maintain their feeding site, these secondary pathogens can cause vine decline and collapse. The resulting exposure of fruit to sunscald compounds further reduces marketable yield.

Previous research has explored alternative management strategies, including planting resistant pepper cultivars (Djian-Caporalino et al., 2011) and, in some cases, deploying susceptible tomato varieties to manage resistance-breaking populations (Castagnone-Sereno et al., 2007). These studies found that resistance breaking imposes reproductive fitness costs in some populations but not others. Nevertheless, susceptible cultivars can serve as a useful stopgap to manage resistance-breaking populations until alternative sources of resistance becomes available (Parlevliet, 1979; Pariaud et al., 2009; Mundt, 2014). However, these studies did not use a diverse set of resistance-breaking populations from a defined geographic region, and they evaluated a limited number of crop hosts that are rarely grown in standard rotation with processing tomatoes. Relying on such hosts would force growers to alter their cultivation plans and purchase new equipment, making them impractical as a sustainable long-term solution.

Although crop rotation can substantially reduce pathogen pressure when non-host or poor-host crops are planted, RKNs are notorious for their broad host range, which leaves growers with few effective rotation options. Current rotation crops for tomatoes in California include safflower, sunflower, cotton, cucurbits, corn, alfalfa, wheat, and dry beans, all of which can be susceptible to RKN infection. The distribution, species identity, and genetic diversity of resistance-breaking RKNs across California remains poorly characterized, limiting the development of practical rotation-based management strategies. In this study, we collected 14 resistance-breaking RKN isolates, established single egg mass isolates, determined their species identity, and compared their genomes to investigate genetic diversity. We then evaluated nematode reproduction on four crops currently used in rotation with processing tomatoes, along with susceptible tomato, in the absence of *Mi-1*. Alfalfa supported very low nematode reproduction across all isolates tested, whereas reproduction on the other rotation crops varied among isolates. These findings highlight the need to tailor crop rotation strategies to the RKN population present in each field.

## Methods

### Plant Material, Germination, and Growth

Five plant species were used in this study: tomato (*Solanum lycopersicum* cv. Moneymaker), cucumber (*Cucumis sativus* cv. Marketmore76), alfalfa (*Medicago sativa* cv. Ameristand 433 TRR), corn (*Zea mays* cv. Passion II), and wheat (*Triticum aestivum* cv. Keldin). Surface-sterilized tomato and alfalfa seeds were germinated on moist filter paper in the dark at 28°C, and seedlings were transplanted after 7 days. Cucumber, corn, and wheat seeds were sown directly into 1-L pots containing sterile, free-draining coarse sand. Plants used for infection assays and nematode cultures were maintained in a greenhouse at a maximum temperature of 26°C and received nutrient irrigation for 3 min three times daily.

### Nematode Isolation and Establishment of Single Egg Mass Cultures

Nematode isolates were collected from 14 independent processing tomato field sites in California. Fields exhibiting patches suggestive of RKN infection were identified by the University of California Cooperative Extension (UCCE) advisors, and sampling was arranged accordingly. Major soil series was recorded for each location site by using the United States Department of Agriculture Natural Resources Conservation Service (USDA-NRC) on the soil surveys platform (websoilsurvey.nrcs.usda.gov/app/WebSoilSurvey.aspx). GPS coordinates were used to identify collection locations and provided soil maps helped identify the major soil types.

Whole roots were collected from the border regions of patches within each field. Between 3 and 8 root samples were collected from each field site, and placed in 3.8-L Ziplock bag, and kept in a cooler until they were brought to the lab. GPS coordinates were recorded for each sample site. The distribution map was generated in Google Maps and assembled in Microsoft PowerPoint. Physical distance for between each sampling site was generated using omni calculator latitude/longitude distance calculator (www.omnicalculator.com/other/latitude-longitude-distance). Root samples from each location were pooled, and nematode eggs were extracted as described previously (Blundell et al., 2026). Eggs were counted and 15,000 eggs were inoculated onto susceptible (cv. MV) and resistant tomato (cv. VFNT, which is homozygous for *Mi-1*) (Wang et al., 2009), five plants each, to confirm resistance-breaking. After 5 weeks, roots were rinsed with water to remove the sand. Roots were then evaluated for gall presence, then stained with Erioglaucine (50 µg/mL) (Sigma-Aldrich, cat. 861146) to view egg mass presence. Positively infected roots on resistant tomato were used to isolate 8 single egg masses. In a 24-well plate, 1 mL of sterile water and an individual egg mass was added to each well. A total of 8 egg masses were collected for each sample location. Hatched J2s from individual wells were inoculated on resistant tomato, each plant representing a single egg mass. After 6 months of culturing, roots were harvested and eggs were counted. For each field site, the single-egg mass (SEM) culture with the highest total egg count was used for further analysis.

### Nematode Perineal Patterns

Adult female RKNs were extracted from purified culture plants ∼3 months after infection on resistant tomato (cv. VFNT). Sand was rinsed from the roots and females were picked individually from galled tissues. Individual females were placed on a microscope slide and 10 µL of 45% lactic acid was added. At least 5 female nematodes were prepared and visualized for each isolate. The females were cut in half, and the internal contents were dissolved. The lower half of the nematode was positioned towards the slide, and the lactic acid was removed replacing it with 15 µL of glycerol to prevent drying. A coverslip was placed to flatten the nematode posterior against the slide. Prepared slides were sealed with clear nail polish and examined for patterns and clarity using a compound microscope (Leica DM1700, Leica Microsystems, Wetzlar, Germany) and images were taken using LAS X Scientific Software. Perineal patterns were compared with Eisenback et al., 1981 for reference and species identification.

### DNA Extraction

Nematode eggs were collected and surface-sterilized as described in Nematode Fitness assays and Culture Maintenance section. After sterilization, eggs were equally proportioned into two 2-mL Eppendorf tubes and spun at 6,000 RPM for 1 minute. The supernatant was discarded, and nematode eggs were flash-frozen in liquid nitrogen. Samples were stored at -80^°^C until DNA extraction. DNA Extraction was adapted from Godinez-Vidal and Groen, 2025. Frozen eggs were placed in a pre-cooled mortar and pestle along with 200 µL of Sodium Dodecyl Sulfate – Extraction Buffer (SDS-EB) and ground to a fine powder. Ground samples were returned to a 1.5 mL tube and an additional 300 µL of SDS-EB buffer and 100 µL of Proteinase K were added. Preheated cetyltrimethylammonium bromide (CTAB) buffer (500 µL, 65°C) was added to the sample and inverted. Samples were incubated at 60^°^C for 30 minutes gently inverting every 5 minutes. The sample was transferred to a 2 mL tube and 1 mL of phenol: chloroform was added and inverted for 2 minutes. Samples were centrifuged for 5 minutes at 13,000 RPM. The supernatant was transferred to a new 1.5 mL tube. RNase A (10 µL, 20mg/mL) was added to the sample and incubated for 15 minutes at 37^°^C. Chloroform-isoamyl alcohol (24:1) 1 mL was added and the sample was inverted for 2 minutes following by centrifugation as before. The supernatant was transferred to a new tube. Afterwards, 5 M Ammonium Acetate (NH4OAc) was mixed into the supernatant for a final concentration of 0.75 M. An equal volume of isopropanol was added to the total volume, and the tubes were mixed by inversion (Dai et al., 2026). The tubes were incubated at -80^°^C for 30 minutes and centrifuged the same as before to discard the supernatant. Next, 1 mL of 70% ethanol was added to wash the pellet, followed by centrifugation at 13,000 RPM for 1 minute to remove the supernatant. The samples were spun down again and the pellet air dried for 3 more minutes at room temperature. The DNA pellet was dissolved in 30 µL of sterile water. DNA concentrations were taken on a NanoDrop (Thermo Fisher Scientific, cat. 250577).

### Mitochondrial Sequence-Based Species Identification

Extracted DNA was adjusted to 100 ng/µL for each PCR reaction. New England Biolabs Taq Polymerase and Buffer were used according to the manufacturer’s protocol (cat. M0273S). The mitochondrial primer set MORF (5’ –ATC GGG GTT TAA TAA TGG G – 3’) and MTHIS (5’ – AAA TTC AAT TGA AAT TAA TAG C – 3’) was used for PCR reactions at an annealing temperature of 50^°^C (Pagan et al., 2015). The PCR products were purified using a Zymo DNA Clean & Concentrator-25 kit (cat. D4033), and the concentrations were measured on a NanoDrop spectrophotometer (Thermo Scientific, cat. 250577). Following purification, DNA sample concentrations were adjusted to 100 ng/10 µL for Sanger Sequencing (Quintara Biosciences, Cambridge, MA). Sequences were analyzed using NCBI-Bn and in WormBase ParaSite (parasite.wormbase.org).

### Genome Sequencing and PCA Analysis of Single Nucleotide Polymorphisms

Principal Component Analysis (PCA) was performed to examine genetic variation among the collected isolates and to help visualize population structure. DNA prepared from SEM isolates was submitted to Novogene Corporation Inc. (Sacramento, CA, USA for Illumina sequencing). Samples were checked for quality, and Illumina sequencing was performed on a NovaSeq X Plus instrument (Novogene). Single nucleotide polymorphisms (SNPs) were identified and mapped to the reference genomes for *M. javanica*, *M. incognita*, and *M. arenaria* (Dai et al., 2023). Illumina paired-end reads were aligned to the reference genome using bwa-mem (Li & Durbin 2009). SNPs were identified using the Genome Analysis Toolkit (GATK, v4.6.1.0) with HaplotypeCaller (McKenna et al., 2010). Raw variants were filtered according to standard quality metrics, and only high-confidence SNPs were retained for downstream analyses. Principal component analysis (PCA) was performed in R (v.4.5.2). VCF files were imported and processed in R (v4.5.2) using vcfR package (v1.15.0). PCA was performed based on the genotype matrix derived from the filtered high-confidence SNPs, with PC1 and PC2 representing the first and second orthogonal axes explaining the largest proportions of genetic variance among isolates, respectively. PCA plots were generated using ggplot2 (v3.5.1).

### Nematode Fitness Assays and Culture Maintenance

Nematode egg collection and surface sterilization were conducted as described in Yimer et al., 2023 on resistant tomato cv. VFNT. *M. incognita eggs* were hatched as described in Blundell et al., 2026. For each plant, 500 hatched J2s in 1mL water were inoculated. Plants tested for reproduction included susceptible tomato (cv. Moneymaker), cucumber (cv. Marketmore76), alfalfa (cv. Ameristand 433 TRR), corn (cv. Passion II), and wheat (cv. Keldin). Ten plants per nematode isolate were inoculated per experiment, with three replicate experiments per crop. Plants were harvested for egg collection at 35 days post inoculation (dpi). Intact roots were shaken in a 10% bleach (6% NaClO) solution for 3 minutes and rinsed over a set of sieves (75 µm and 25 µm) with water. Eggs were collected from the 25 µm sieve into a 50 mL Falcon tube. Eggs were counted for all crops tested.

### Statistical Analysis

For all infection assays, GraphPad Prism (https://www.graphpad.com/) was utilized for statistical analysis. Outliers were removed using Prism’s ROUT method with Q = 10% and cleaned data were used for one-way ANOVA with a 95% confidence interval, assuming a Gaussian distribution.

### Data Availability

The raw Illumina sequencing reads generated for the resistance-breaking isolates in this study have been deposited in the NCBI Sequence Read Archive (SRA) under BioProject accession PRJNA1545056 (https://www.ncbi.nlm.nih.gov/bioproject/PRJNA1545056).

## Results

### RKN Populations from All 14 Field Collections Were Resistance-Breaking

To characterize resistance-breaking RKNs associated with symptomatic processing tomato fields, we collected RKN populations from 14 fields planted with tomato cultivars carrying *Mi-1*. Fields with distinct patches of diseased plants were selected, and GPS coordinates were recorded for each collection site (**Figure 1**, **Table 1**, and **Supp. Table 1)**. All sampled fields were located in California’s Central Valley, the major processing tomato production region of the state. Fields were sampled between June and October 2025, when plants were at a late growth stage.

**Figure 1.**
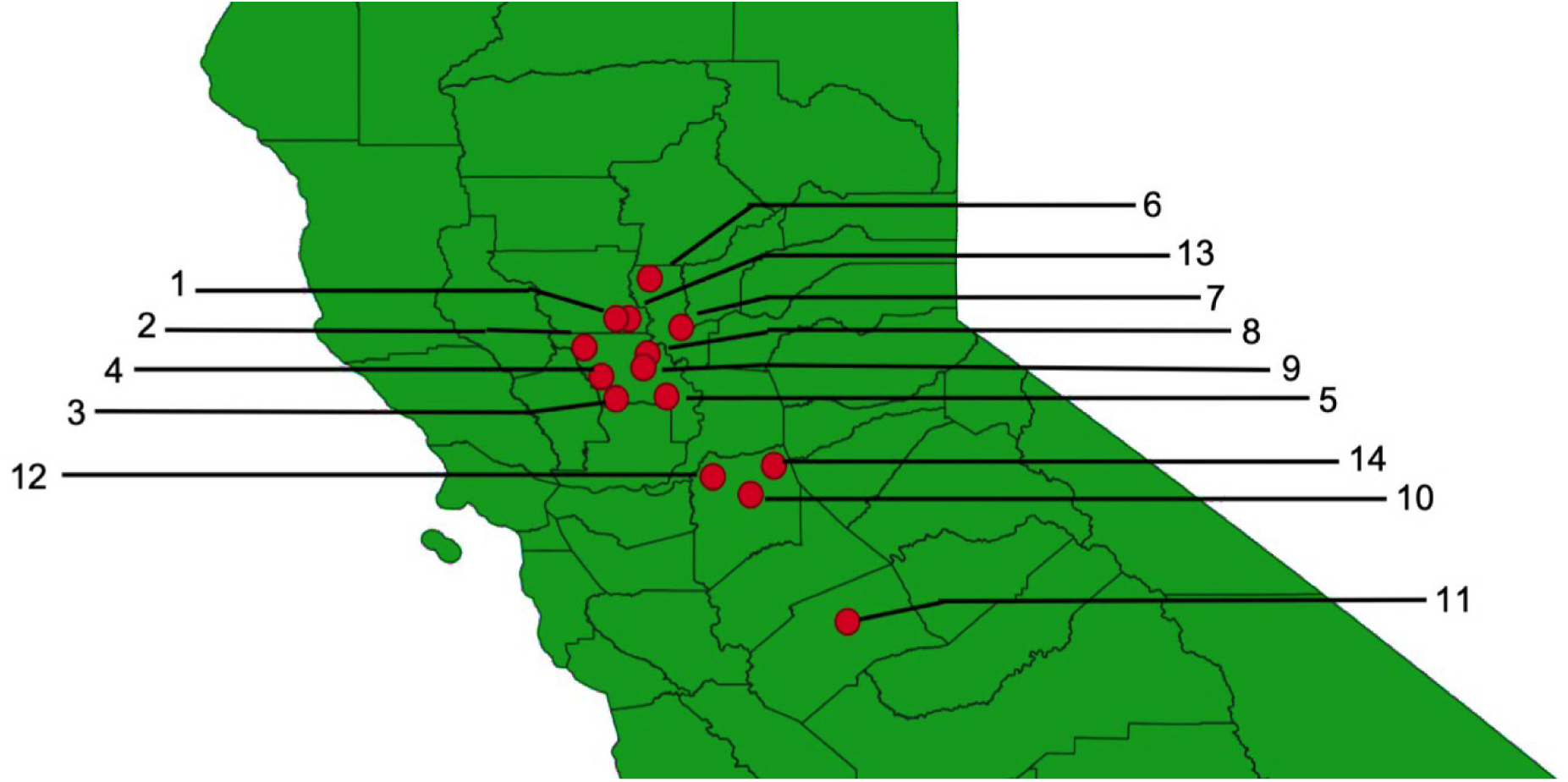
Collection sites of resistance-breaking RKN isolates in California processing tomato fields. Isolates were collected from processing tomato fields in California’s Central Valley, USA. Each collection site is marked with a red circle and labeled with the isolate number.

**Table 1.** Collected resistance-breaking isolates of RKN species identification and locations. Isolate number, California County, soil type, isolate name and species identification are depicted for each isolate.

| **Isolate** | **California County** | **Soil Type** | **Strain Name** | **Species Identification** |
|---|---|---|---|---|
| 1 | Solano County | Silty Clay Loam | AB1 | *Meloidogyne incognita* |
| 2 | Yolo County | Silt Loam | AB2 | *Meloidogyne incognita* |
| 3 | Yolo County | Loam | AB3 | *Meloidogyne incognita* |
| 4 | Yolo County | Silt Clay Loam | AB4 | *Meloidogyne incognita* |
| 5 | Yolo County | Silt Loam | AB5 | *Meloidogyne incognita* |
| 6 | Sutter County | Silt Loam, Clay | AB6 | *Meloidogyne incognita* |
| 7 | Sutter County | Sandy Loam | AB7 | *Meloidogyne incognita* |
| 8 | Yolo County | Sandy Loam | AB8 | *Meloidogyne javanica* |
| 9 | Yolo County | Sandy Loam | AB9 | *Meloidogyne javanica* |
| 10 | San Joaquin County | Clay | AB10 | *Meloidogyne javanica* |
| 11 | Merced County | Sandy Loam | AB11 | *Meloidogyne incognita* |
| 12 | San Joaquin County | Muck | AB12 | *Meloidogyne incognita* |
| 13 | Solano County | Loam | AB13 | *Meloidogyne incognita* |
| 14 | San Joaquin County | Clay Loam | AB14 | *Meloidogyne arenaria* |

To determine whether collected populations contained resistance-breaking nematodes, we conducted greenhouse infection assays on susceptible (cv. MV) and resistant (homozygous *Mi-1* carrying; cv. VFNT) tomato at a maximum temperature of 26^°^C. Successful infection was scored by the presence of galls and egg masses on the roots. All 14 field populations produced galls and egg masses on both susceptible and resistant tomato, confirming the presence of resistance-breaking nematodes at every sampled site. Single egg mass cultures (SEM) were established from egg masses collected from resistant tomato roots. One culture from each field population was selected for subsequent analyses, yielding 14 isolates designated AB1–AB14.

To identify SEM isolates to species, genomic DNA was amplified using mitochondrial primers MORF/MTHIS and amplicons were subjected to Sanger sequencing (Pagan et al., 2015; **Supp. Table 2**). Of the 14 SEM isolates, ten were identified as *M. incognita*, three as *M. javanica*, and one as *M. arenaria*. To corroborate the molecular identifications with morphology, perineal patterns were obtained for all 14 isolates and compared against Eisenback et al. (1981). A representative pattern for each of the three MIG species is shown in **Figure 2**, and patterns for all populations are compiled in **Supp. Figure 1**. Because MIG species share overlapping morphological characters, identifications relied on subtle differences in perineal patterns. *M. incognita* (AB1, AB2, AB3, AB4, AB5, AB6, AB7, AB11, AB12, and AB13) was distinguished by a high, squarish dorsal arch and the absence of lateral folds (**Figure 2; Supp. Figure 1**). *M. javanica* (AB8, AB9, AB10) was identified by a subdued, flattened dorsal arch and the presence of lateral folds (**Figure 2; Supp. Figure 1**). *M. arenaria* showed a dorsal arch with shoulders and striae near the lateral lines (**Figure 2; Supp. Figure 1**).

**Figure 2.**
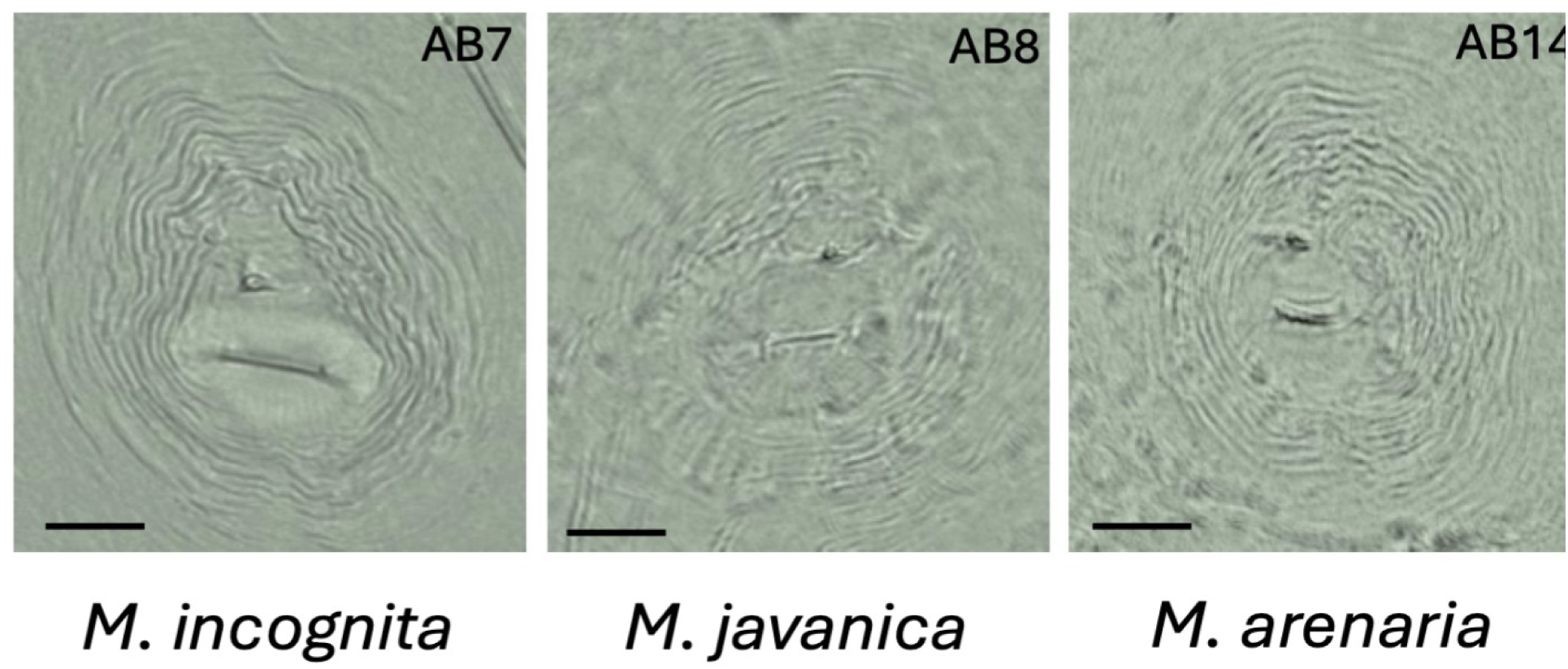
Representative perineal patterns of the three MIG-group *Meloidogyne* species. Patterns are shown for *M. incognita* (AB7), *M. javanica* (AB8), and *M. arenaria* (AB14). Scale bars = 25 µm.

### Resistance-Breaking *Meloidogyne incognita* Isolates Are Genetically Diverse

To characterize genetic diversity among the collected RKN populations, we sequenced the genomes of the SEM isolates using Illumina platform. High-quality single-nucleotide polymorphisms (SNPs) identified relative to *M. incognita* reference genome (Dai et al., 2023) were used for principal component analysis (PCA). Given the predominance of *M. incognita* among the collected isolates, we focused the PCA on genetic variation within this species (**Figure 3**). PC1 and PC2 explained 19.27% and 18.08% of the variance, respectively.

**Figure 3.**
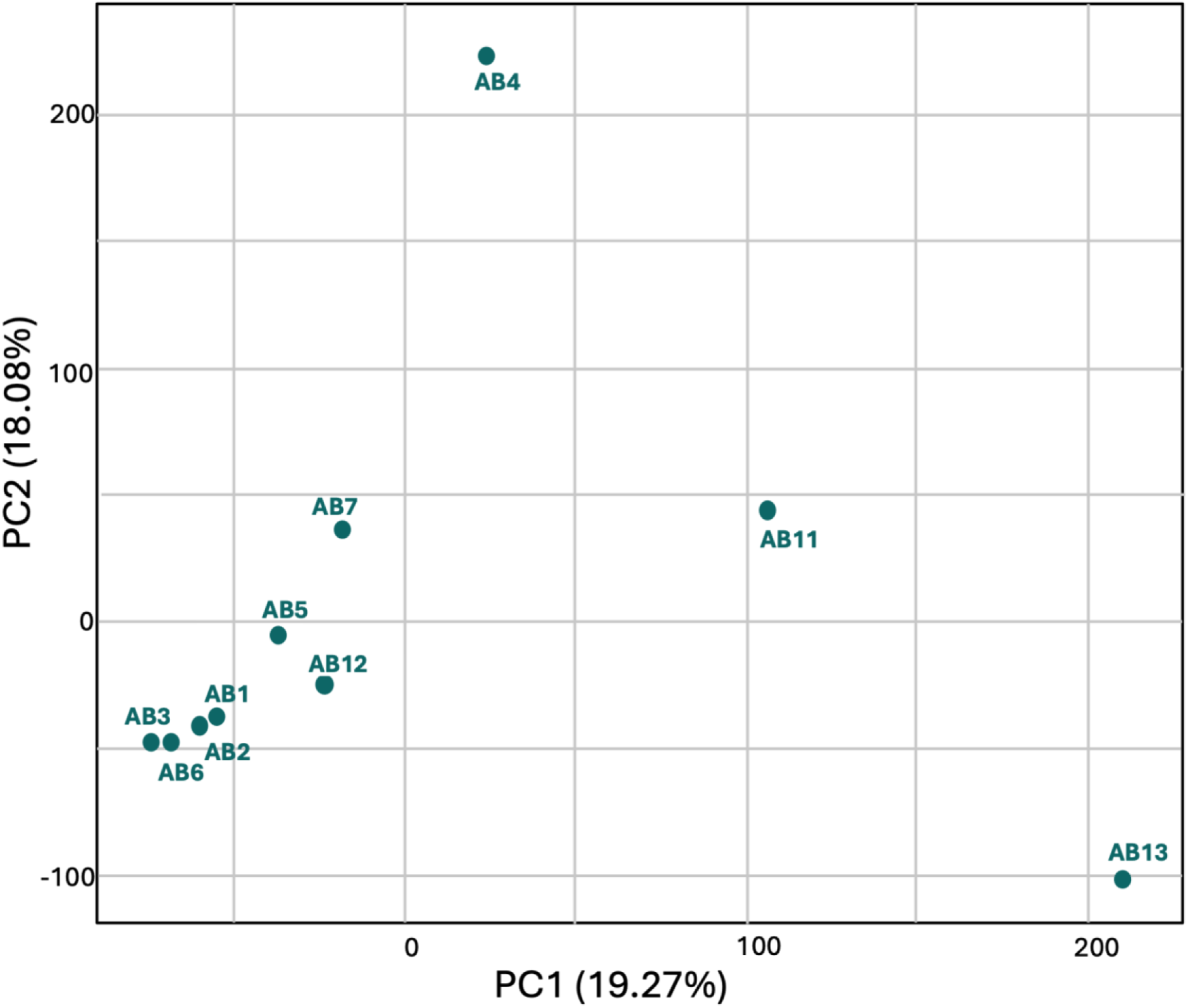
Principal component analysis (PCA) of the collected *Meloidogyne incognita* isolates. The PCA was generated from SNPs called against the *M. incognita* reference genome (WHF4-1, China). PC1 and PC2 explain 19.27% and 18.08% of the variance, respectively. Isolates positioned closer together are more genetically similar, whereas those positioned farther apart are more divergent.

AB1, AB2, AB3, and AB6 clustered tightly along both axes, indicating close genetic similarity, while AB5, AB7, and AB12 were positioned near this cluster. In contrast, AB4 was separated from the main cluster primarily along PC2, whereas AB13 was clearly separated along PC1. AB11 occupied an intermediate position between the main cluster and the divergent isolates AB4 and AB13. Overall, these results indicate that resistance-breaking *M. incognita* occurs in multiple genetic backgrounds. This pattern is consistent with independent emergence of resistance breaking at multiple locations, potentially followed by local spread, but does not exclude dissemination and subsequent divergence of a shared ancestral population.

### Reproduction on Rotation Crops Varied Among Resistance-Breaking Isolates

Based on their position in PCA, four most divergent isolates (AB1, AB4, AB11, and AB13) were selected to evaluate reproduction on crops currently growin in rotation with processing tomatoes. Four rotation crops were tested: alfalfa (cv. Ameristand 433 TRR), corn (cv. Passion II), cucumber (cv. Marketmore76), and wheat (cv. Keldin) along with susceptible tomato (cv. Moneymaker). These crops are used in rotation with processing tomato and have been reported as hosts or potential hosts of *M. incognita*.

Nematode reproduction was assessed by counting eggs at 35 days post-inoculation (dpi). Egg production was used because it was previously shown to be the reproductive parameter most affected when a resistance-breaking isolate was tested on susceptible cultivars (Blundell et al., 2026). *M. incognita* isolate VW6, which does not reproduce on tomato carrying *Mi-1*, was included as a reference for reproductive potential on each host.

On susceptible tomato, egg production was similar to that of VW6 for most isolates, except for AB1, which produced significantly fewer aggs (**Figure 4A**). Cucumber supported higer reproduction by all isolates, with AB1 and AB11 producing significantly more eggs than VW6 (**Figure 4B**). Alfalfa supported the lowest reproduction by both VW6 and the resistance-breaking isolates, with egg counts near zero in all cases (**Figure 4C**).On corn, AB1 and AB11 also reproduced significantly more eggs than VW6, whereas AB4 and AB13 reproduced poorly, indicating that corn supported some resistance-breaking isolates more than others (**Figure 4D**). On wheat, AB1, AB4, and AB13 produced significantly fewer eggs than VW6, whereas AB11 did not differ significantly from VW6 (**Figure 4E**). These data show that reproduction on rotation crops varied by isolate, indicating that host suitability was population dependent.

**Figure 4.**
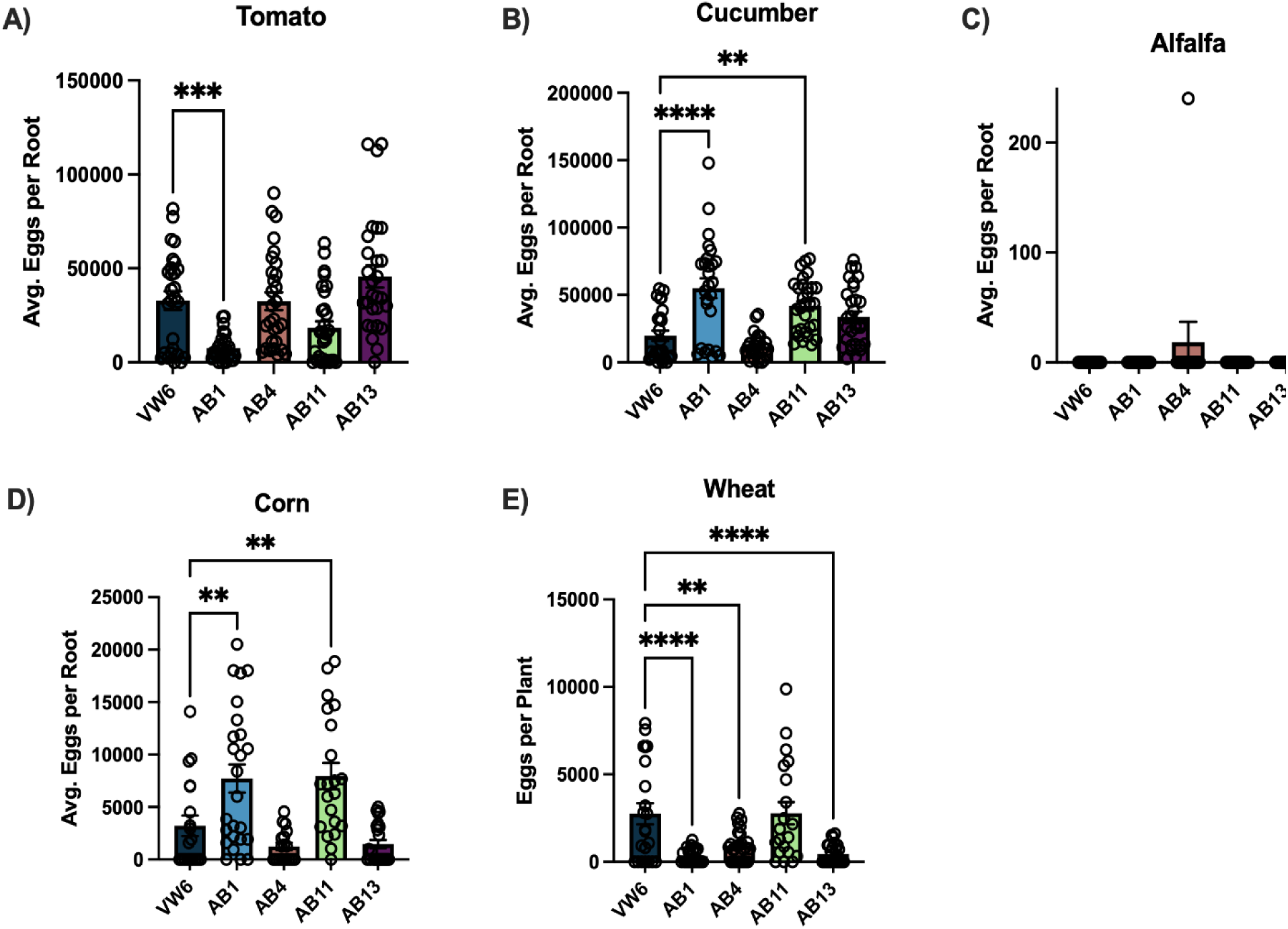
Infection assays evaluating nematode virulence on rotation crops showed alfalfa to suppress nematode reproduction. Reproduction was evaluated at 35 dpi on each host. The rotation crops cucumber (cv. Marketmore76), alfalfa (cv. Ameristand 433 TRR), corn (cv. Passion II), and wheat (cv. Keldin), along with susceptible tomato (cv. Moneymaker), were inoculated with the four most divergent *M. incognita* isolates collected; the wild-type VW6 was included as a control for comparison on each crop. Each data point represents an individual root system, and bars indicate the mean ± standard error of the mean (SEM). Outliers were removed using the ROUT method (Q = 10%), and an ordinary one-way ANOVA with Dunnett’s multiple comparisons test was used to compare each isolate to VW6. Asterisks indicate significance (p < 0.05).

This pattern was reinforced when the data was reorganized to evaluate each isolate separately across crops (**Figure 5A–E**). AB1 reproduced least on alfalfa, followed by wheat and then susceptible tomato. AB4 and AB13 likewise reproduced least on alfalfa, with wheat and corn also supporting low reproduction. For AB11, reproduction was lowest on alfalfa, followed by corn. Overall, all isolates reproduced abundantly on cucumber and poorly on alfalfa, indicating that tested cucumber cultivar was a favorable host for these resistance-breaking isolates, whereas the tested alfalfa cultivar was consistently a poor host.

**Figure 5.**
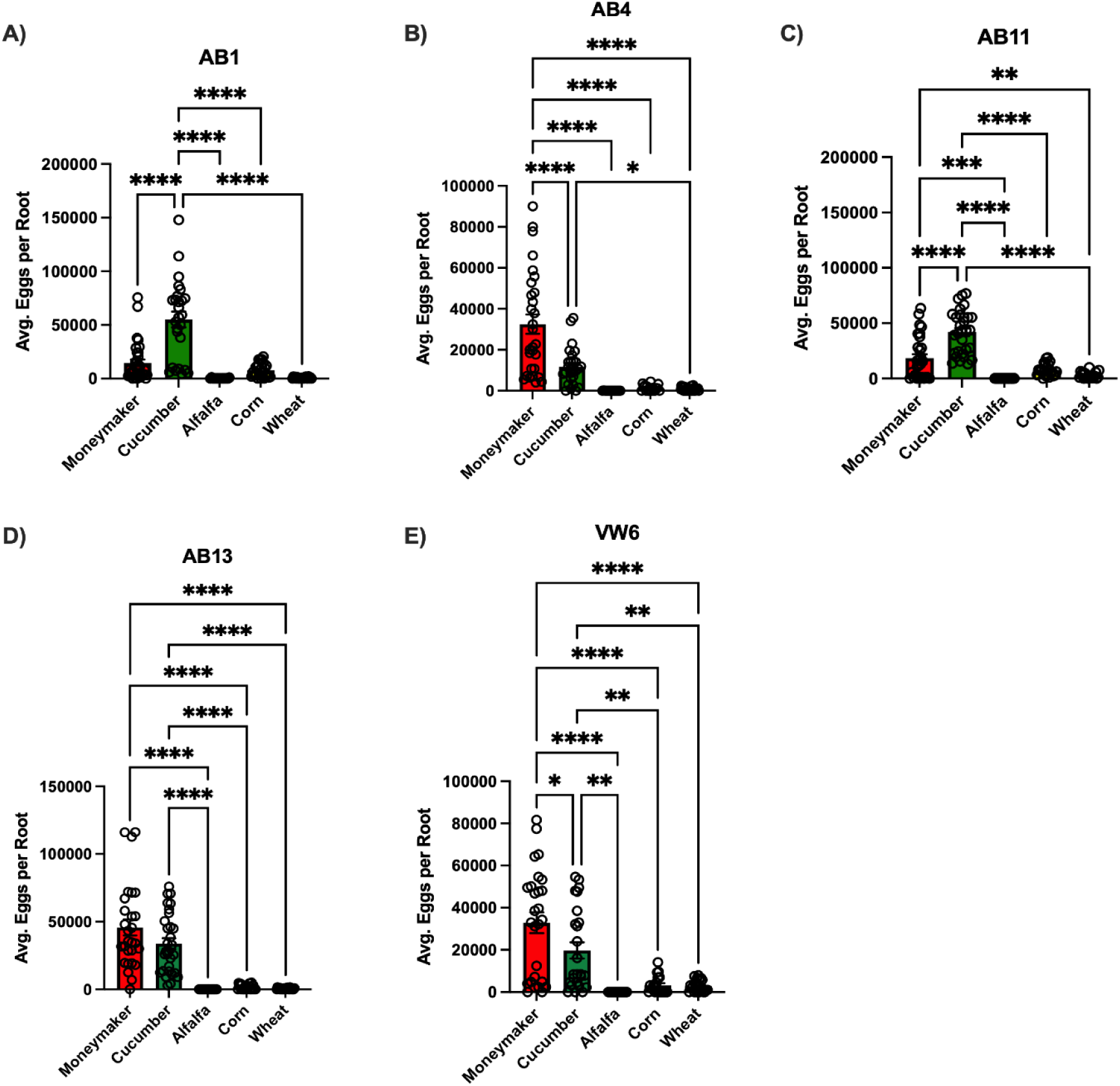
Infection assays on rotation crops showed reproductive differences among individual resistance-breaking *M. incognita* isolates. Infection assays were conducted on susceptible tomato (cv. Moneymaker), cucumber (cv. Marketmore76), alfalfa (cv. Ameristand 433 TRR), corn (cv. Passion II), and wheat (cv. Keldin) for four resistance-breaking isolates and one wild type (VW6). Reproduction on each host is shown as the number of eggs produced at 35 dpi. Each data point represents an individual root system, and bars indicate the mean ± standard error of the mean (SEM). Outliers were removed using the ROUT method (Q = 10%), and an ordinary one-way ANOVA with multiple comparisons was conducted. Asterisks indicate significance (p < 0.05).

## Discussion

In this study, we collected resistance-breaking root-knot nematodes from fourteen tomato fields in California and evaluated their reproduction on crops commonly used in processing tomato rotations. Morphological and genomic analyses identified ten isolates as *M. incognita*, three as *M. javanica,* and one as *M. arenaria* (**Figure 2** and **Supp. Table 1**). A SNP-based PCA of the *M. incognita* isolates, generated through Illumina whole-genome sequencing, revealed substantial genetic diversity among *M. incognita* isolates (**Figure 3**). This diversity suggests that resistance-breaking may have evolved independently at multiple locations, potentially in response to repeated cultivation of *Mi-1*-carrying tomatoes, followed by local spread in some cases. The close genetic similarity among AB1, AB2, AB3, and AB6 is also consistent with dissemination among fields. In particular, AB1, AB2, and AB3 were collected from geographically close sites, supporting the possibility of local spread among neighboring fields (**Supplementary Table 1**). Nevertheless, SNP-based PCA alone is not sufficient to establish independent origins of resistance-breaking or reconstruct its spread.

Among the four tested isolates, alfalfa consistently supported very low reproduction, whereas cucumber supported abundant reproduction across all isolates tested (**Figure 4**). These results identify the tested alfalfa cultivar as a poor host for all four isolates and support its potential use in rotation-based nematode management. However, environmental conditions may not have been optimal for RKN infection on alfalfa, as it is typically grown in cooler climates (21^°^C – 24^°^C), and other cultivars may differ in host susceptibility. Its broader effectiveness therefore requires evaluation across cultivars and under field conditions. In contrast, tested cucumber cultivar appeared to be a consistently favorable host, supporting high levels of reproduction across all tested isolates. Planting cucumber after tomato could therefore sustain or increase resident *M. incognita* populations rather than suppress them, making it an unsuitable rotation choice for reducing these populations. Other cucurbits, such as zucchini or squash, may pose a similar risks, but their host suitability should be evaluated directly for each crop and cultivar.

Responses on corn and wheat were particularly informative because reproduction varied among isolates, indicating that these isolates can support some resistance-breaking populations more readily than others. However, it is also plausible that other corn and wheat cultivars may support lower reproduction, although this possibility requires direct evaluation. Consequently, suitability of corn and wheat for rotation-based nematode management may depend on both the nematode population and the cultivar planted. This finding has notable practical implications as corn and wheat are established rotation options in California processing tomato systems, yet their management value may vary with RKN population present in the field. Together, these results support tailoring rotation strategies to local nematode populations, with the tested alfalfa cultivar emerging as a promising option across the isolates evaluated.

Several limitations should be considered when interpreting these results. First, it remains unclear whether resistance-breaking is more frequent in *M. incognita* than other RKN species or whether *M. incognita* is simply more abundant in the sampled region, thereby increasing its representation among our isolates. A previous study of resistance-breaking isolates in California also identified *M. incognita* as most prevalent (Ploeg et al., 2023). However, more intensive and systematic sampling is needed to establish whether this pattern reflects species-level differences in resistance-breaking potential or regional species prevalence. Second, mixed RKN populations may have been present at individual field sites. Because we established single-egg-mass cultures from each population, co-occurring species or genotypes, including *M. javanica* or *M. arenaria*, may have been excluded from downstream assays. Third, plants at all sampled sites showed symptoms suggestive of secondary fungal infection in addition to RKN damage, but the suspected fungal pathogens were not identified. Root damage caused by RKNs may increase the risk of secondary infection by wounding roots and creating entry points for pathogens, but this relationship was not directly tested here and warrants future investigations. Finally, Finally, sampling was limited to 14 fields, restricting the generalizability of our findings. Broader, systematic surveys are needed to characterize the distribution, frequency, and diversity of resistance-breaking RKN populations across California.

Despite these limitations, this study integrates genomic and phenotypic analyses to characterize resistance-breaking RKNs from California processing tomato fields. The observed genetic diversity and variation in reproduction among isolates highlight the importance of considering local nematode populations when designing management strategies. Practically, tested alfalfa cultivar emerged as a promising rotation option across the isolates evaluated, whereas the suitability of other crops varied among isolates. Beyond these findings, the genomically characterized isolates generated here provide a foundation for investigating the genetic basis of resistance-breaking and for developing diagnostic tools that enable rapid screening and targeted management. Overall, durable management of resistance-breaking RKNs will likely require an integrated strategy that combines resistant cultivars, avoidance of highly suitable hosts, and selection of poor-host rotation crops guided by the local RKN population.

## Funding and Disclaimer

This work was funded by the California Department of Pesticide Regulation (24-PMG-R001). The contents may not necessarily reflect the official views or policies of the State of California.

## Acknowledgments

We would like to thank the University of Montana, specifically the Dyer Lab for the wheat seeds, and AgSeeds Unlimited for providing corn and alfalfa seeds.

## Supplemental Data

**Supplementary Table 1.** Distance between each field site sample in kilometers (km). GPS coordinates were computed against each other to acquire distances between each field sampling site by inputting points both latitude and longitude. Distance was generated in omni calculator latitude/longitude distance calculator.

|  | **AB1** | **AB2** | **AB3** | **AB4** | **AB5** | **AB6** | **AB7** | **AB8** | **AB9** | **AB10** | **AB11** | **AB12** | **AB13** | **AB14** |
|---|---|---|---|---|---|---|---|---|---|---|---|---|---|---|
| **AB1** |  | 5.20 | 8.38 | 11.00 | 32.91 | 37.39 | 58.22 | 14.58 | 14.51 | 88.03 | 176.70 | 51.90 | 5.72 | 80.46 |
| **AB2** | 5.20 |  | 10.08 | 7.72 | 28.14 | 33.57 | 53.54 | 11.25 | 11.02 | 93.04 | 181.80 | 56.90 | 10.85 | 85.20 |
| **AB3** | 8.38 | 10.08 |  | 9.06 | 31.25 | 33.05 | 55.70 | 11.40 | 11.64 | 85.70 | 174.72 | 49.69 | 8.78 | 76.87 |
| **AB4** | 11.00 | 7.72 | 9.06 |  | 22.67 | 26.39 | 47.66 | 3.62 | 3.51 | 94.77 | 183.80 | 58.75 | 15.21 | 85.80 |
| **AB5** | 32.91 | 28.14 | 31.25 | 22.67 |  | 12.00 | 25.41 | 19.86 | 19.63 | 115.90 | 204.90 | 80.31 | 37.77 | 105.75 |
| **AB6** | 37.39 | 33.57 | 33.05 | 26.39 | 12.00 |  | 24.29 | 22.82 | 22.88 | 113.40 | 201.90 | 78.86 | 41.12 | 102.03 |
| **AB7** | 58.22 | 53.54 | 55.70 | 47.66 | 25.41 | 24.29 |  | 44.50 | 44.38 | 137.70 | 226.15 | 103.05 | 62.88 | 126.20 |
| **AB8** | 14.58 | 11.25 | 11.40 | 3.62 | 19.86 | 22.82 | 44.50 |  | 0.53 | 96.65 | 185.70 | 60.79 | 18.51 | 87.23 |
| **AB9** | 14.51 | 11.02 | 11.64 | 3.51 | 19.63 | 22.88 | 44.38 | 0.53 |  | 97.00 | 186.05 | 61.12 | 18.57 | 87.63 |
| **AB10** | 88.03 | 93.04 | 85.70 | 94.77 | 115.90 | 113.40 | 137.70 | 96.65 | 97.00 |  | 89.04 | 36.15 | 82.32 | 15.72 |
| **AB11** | 176.70 | 181.80 | 174.72 | 183.80 | 204.90 | 201.90 | 226.15 | 185.70 | 186.05 | 89.04 |  | 125.06 | 171.00 | 100.01 |
| **AB12** | 51.90 | 56.90 | 49.69 | 58.75 | 80.31 | 78.86 | 103.05 | 60.79 | 61.12 | 36.15 | 125.06 |  | 46.2 | 30.05 |
| **AB13** | 5.72 | 10.85 | 8.78 | 15.21 | 37.77 | 41.12 | 62.88 | 18.51 | 18.57 | 82.32 | 171.00 | 46.2 |  | 74.89 |
| **AB14** | 80.46 | 85.20 | 76.87 | 85.80 | 105.75 | 102.03 | 126.20 | 87.23 | 87.63 | 15.72 | 100.01 | 30.05 | 74.89 |  |

**Supplemental Figure 1.**
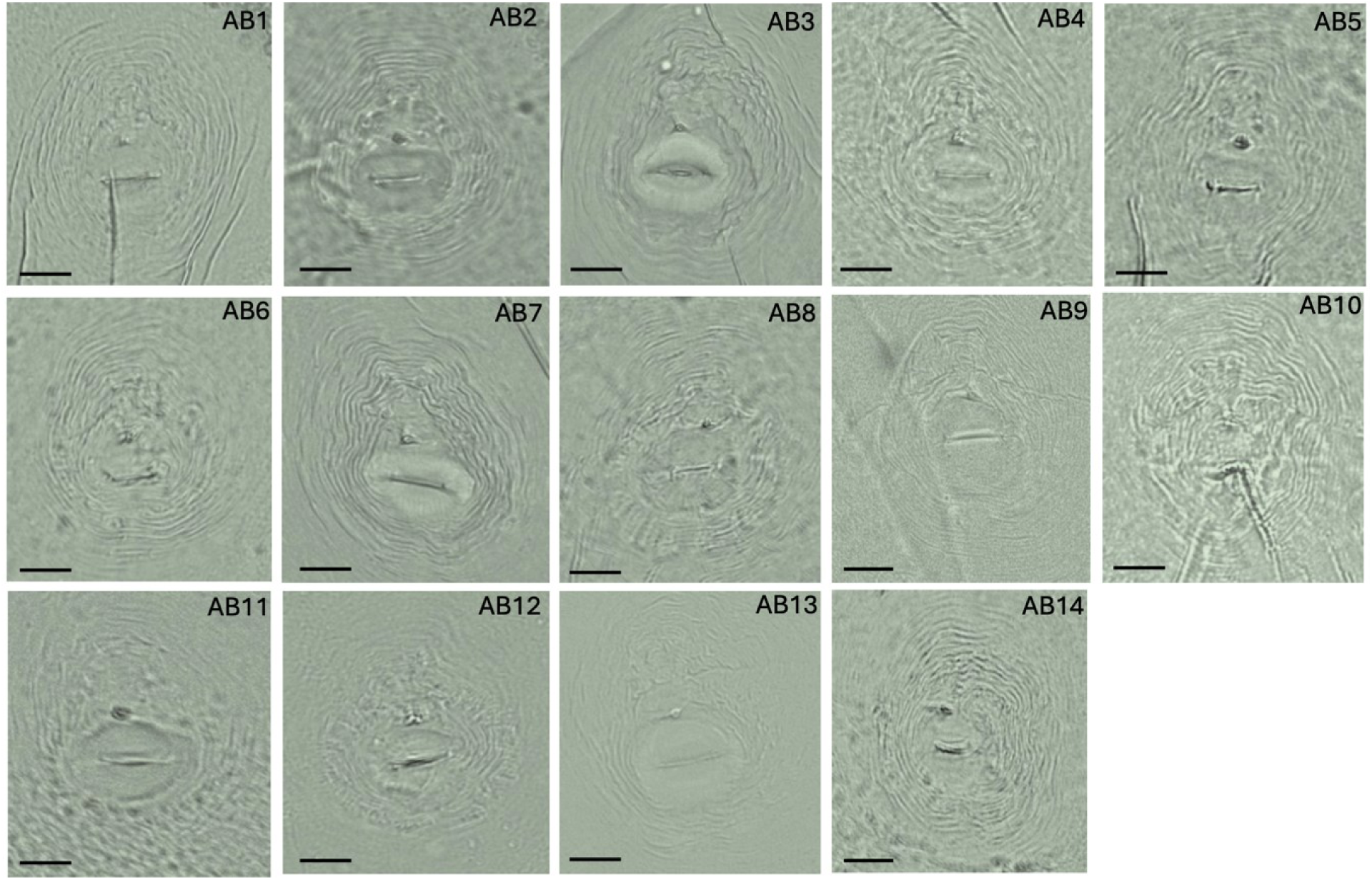
Compilation of all perineal patterns of the fourteen collected isolates. Isolates AB1 – AB14 are listed above corresponding pattern depicted. Light microscopy images are shown under 40x lens. Scale bar = 25 µm.

## Notes

### Competing Interest Statement

The authors have declared no competing interest.

